# CryoPPP: A Large Expert-Labelled Cryo-EM Image Dataset for Machine Learning Protein Particle Picking

**DOI:** 10.1101/2023.02.21.529443

**Authors:** Ashwin Dhakal, Rajan Gyawali, Liguo Wang, Jianlin Cheng

**Author notes:** Corresponding author: Jianlin Cheng. Joint first author (Ashwin Dhakal, Rajan Gyawali).

## Abstract

Cryo-electron microscopy (cryo-EM) is currently the most powerful technique for determining the structures of large protein complexes and assemblies. Picking single-protein particles from cryo-EM micrographs (images) is a key step in reconstructing protein structures. However, the widely used template-based particle picking process is labor-intensive and time-consuming. Though the emerging machine learning-based particle picking can potentially automate the process, its development is severely hindered by lack of large, high-quality, manually labelled training data. Here, we present CryoPPP, a large, diverse, expert-curated cryo-EM image dataset for single protein particle picking and analysis to address this bottleneck. It consists of manually labelled cryo-EM micrographs of 32 non-redundant, representative protein datasets selected from the Electron Microscopy Public Image Archive (EMPIAR). It includes 9,089 diverse, high-resolution micrographs (∼300 cryo-EM images per EMPIAR dataset) in which the coordinates of protein particles were labelled by human experts. The protein particle labelling process was rigorously validated by both 2D particle class validation and 3D density map validation with the gold standard. The dataset is expected to greatly facilitate the development of machine learning and artificial intelligence methods for automated cryo-EM protein particle picking. The dataset and data processing scripts are available at https://github.com/BioinfoMachineLearning/cryoppp

## Background & Summary

Cryo-electron microscopy (cryo-EM) is an experimental technique that captures 2D images of biological molecules and assemblies (protein particles, virus, etc.) at cryogenic temperature using ‘direct’ electron-detection camera technology ^1^. With the advent of cryo-EM, there has been a boom in structural discoveries relating to biomolecules, particularly large protein complexes and assemblies. These 3D structures of proteins ^2^ are important for understanding their biological functions ^3^ and their interactions with ligands ^4,5^, which can aid both basic biological research and structure-based drug discovery ^4,6^. A key step of constructing protein structures form cryo-EM data is to pick protein particles in cryo-EM images (micrographs). Before diving into recent developments in protein particle picking and the bottleneck it faces, it is important to understand the physics and chemistry behind the grid preparation and micrograph image acquisition in cryo-EM experiments.

### I Cryo-EM Grid Preparation and Image Acquisition

The process of acquiring the two-dimensional projections of biomolecular samples (e.g., protein particles) can be summarized in four brief steps: (1) sample purification, (2) cryo-EM grid preparation, (3) grid screening and evaluation, and (4) image capturing. Once the sample is purified according to the standard protocols ^7^; the next step of the single-particle procedure is to prepare the cryo-EM specimen. The grid preparation process, also known as vitrification, is straightforward. An aqueous sample is applied to a grid, which is then made thin. Eventually, the grid is plunged frozen at a time scale that inhibits the crystalline ice formation. Additionally, the particles must be evenly distributed across the grid in a wide range of orientations. It is very difficult to achieve a perfect cryo-EM grid because particles may choose to adhere to the carbon layer instead of being partitioned into holes. They may also adopt preferred orientations within the vitrified ice layer, which reduces the number of unique views ^8^. The grid is ready for analysis once the cryo-EM sample is successfully inserted into the electron microscope ^9^. Images are routinely captured during the screening phase at various magnifications to check for ice and particle quality. After the grids are optimized and ready for cryo-EM data collection, they are taken to a cryo-EM facility where qualified professionals load specimens into the microscope. To enable the best high-quality image capturing, experts adjust several parameters such as magnification, defocus range, electron exposure, and hole targeting techniques (see **Figure 1 (A)-(F)** illustrating the process of preparing cryo-EM samples and acquiring cryo-EM images). More details regarding cryo-EM sample preparation and image acquisition can be found in these studies ^7,10^.

**Figure 1:**
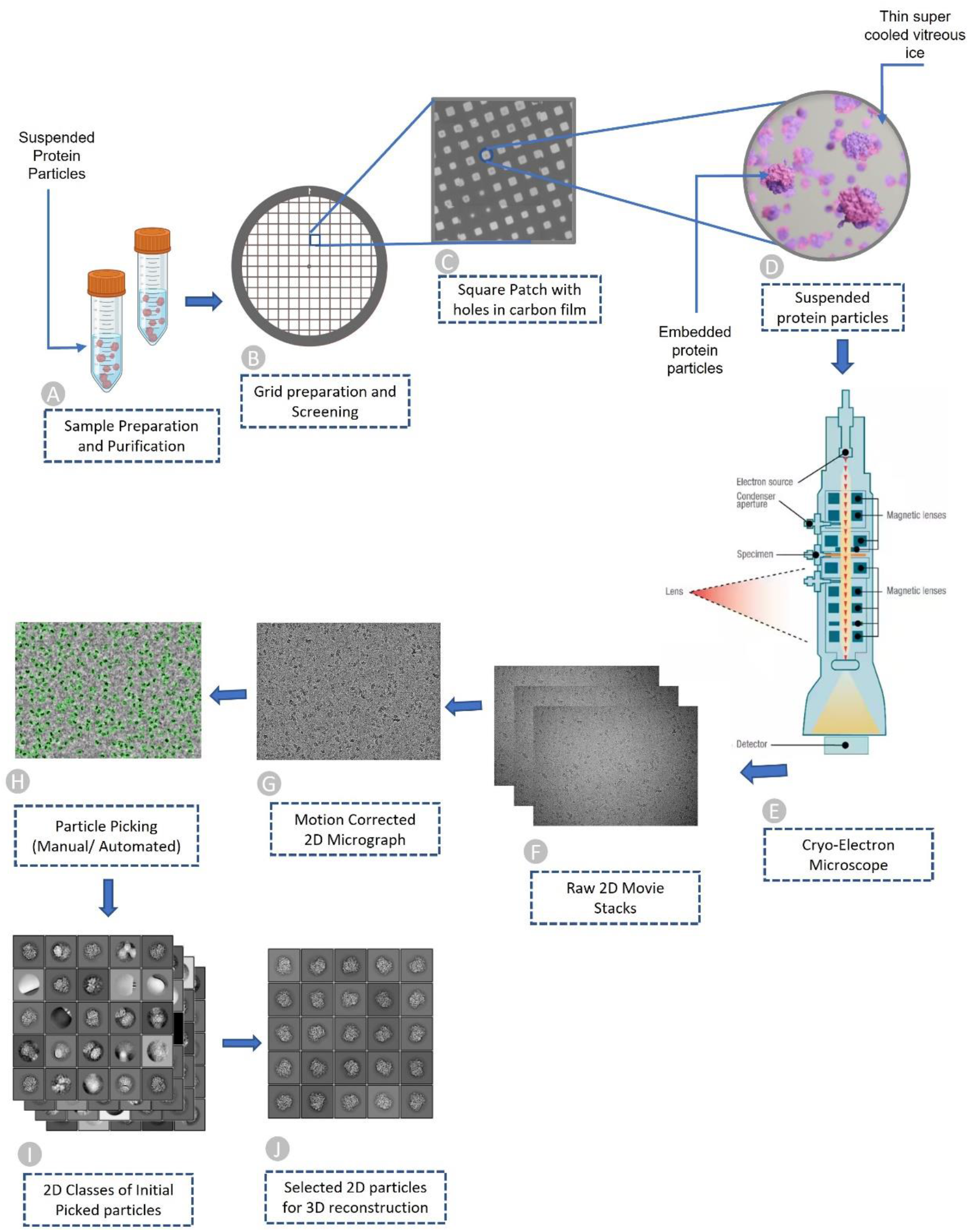
Overview of Cryo-EM pipeline, from sample preparation to particle recognition. (A) Aqueous sample preparation that contains variably dispersed heterogenous structure. (B) Cryo-EM grid containing holes that are filled with dispersed protein particles. (C) Magnified image of square patch illustrating microscopic holes in carbon. (D) Zoomed-in view of single hole containing suspended protein particles in thin layer of vitreous ice. (E) Cryo-Electron microscope used to facilitate high quality image generation. (F) Stack of 2D movie frames generated from microscope, called micrographs. (G) Motion corrected 2D micrograph images. (H) Particle picking using manual intervention or automatic procedures (green circles represent picked particles). (I) Initial 2D classes that contain quality protein particles along with junks and aggregates. (J) Best quality protein particles identified through computational analysis and visual inspection for 3D protein structure reconstruction.

### II Cryo-EM Micrographs and Single Particle Analysis

When the electron beam passes through a thin vitrified sample, it creates 2D image projections (see **Figure 1** for a visual illustration) of the samples (e.g., protein particles). The projections of the particles in various orientations are stored in different image formats (MRC, TIFF, TBZ, EER, PNG, etc.) which are called micrographs. Once the micrographs are obtained, the objective is to locate individual protein particles in each micrograph while avoiding crystalline ice contamination, malformed particles and grayscale background regions. In other words, the input for the particle picking problem is a micrograph, while the desired output is the coordinates of every protein particle in that micrograph (refer to **Figure 1** for the entire pipeline). Accurate detection of particles is necessary, as the presence of false positive particles can complicate subsequent processing, and eventually cause the 3D reconstruction process to fail entirely. The picking task is challenging due to several factors, including high noise levels caused by ice and contamination, low contrast of particle images, and unpredictability in an individual particle’s appearance caused by variation in orientation. Once the particles are extracted from the micrographs, single particle analysis is performed to reconstruct the 3D density map and protein structure.

### III Advances and Challenges in Single Protein Particle Picking

Several research initiatives were carried out worldwide to improve hardware ^11–13^ and software ^14–16^ to streamline and automate the data collection and processing steps for the cryo-EM determination of 3D structures. The recent technological advances in sample preparation, instrumentation, and computation methodologies have enabled the cryo-EM technology to solve massive protein structures at better than 3 A resolution. To obtain a high-resolution protein structure, selecting enough high-quality protein particles in cryo-EM images is critical. However, protein particle picking is still largely a labor-intensive and time-consuming process. One challenge facing cryo-EM data analysis is to develop automated particle picking techniques to circumvent manual intervention. To tackle the problem, numerous automatic and semi-automatic particle-picking procedures have been developed.

A common technique for particle picking, known as template matching, uses user-predefined particles as templates for identifying particles in micrographs through image matching. However, because of varied ice contamination, carbon areas, overlapping particles, and other issues, the template matching often selects invalid particles (e.g., false positives). So subsequent manual particle selection is necessary.

To deal with the issue, artificial intelligence (AI) and machine learning-based approaches have been proposed, which can be less sensitive to impurities and more suitable for large-scale data processing and therefore hold the potential of fully automating the particle picking process. XMIPP ^17^, APPLE picker ^18^, DeepPicker ^19^, DeepEM ^20^, FastParticle Picker ^21^, crYOLO ^22^, PIXER ^23^, PARSED ^24^, WARP ^25^, Topaz ^26^, AutoCryoPicker ^27^, and DeepCryoPicker ^28^, can be taken as good examples of such efforts.

The datasets used to train and test machine learning particle picking methods were curated from EMPIAR ^29^. It contains almost all the publicly available raw cryo-EM micrographs. It is a public repository containing 1,159 entries/datasets (2.39 PB) as of Jan 29, 2023. It includes not just cryo-EM images of proteins, but also Soft X-ray Tomography (SXT), cryo-ET and many other microscopic projections of other biological samples. Only some cryo-EM images of a small number of datasets in EMPIAR contain particles manually labelled by the original authors of the data. Therefore, most existing machine learning methods for particle picking were trained and tested on only a few manually labeled datasets of a few proteins like Apoferritin and Keyhole Limpet Hemocyanin (KLH). The methods trained on the limited amount of particle data of one or a few proteins cannot generalize well to pick particles of various shapes in the cryo-EM micrographs of many diverse proteins in the real world. Therefore, even though machine learning particle picking is a promising direction, no machine learning method has been able to replace the labor-intensive template-based particle picking in practice. Therefore, the lack of manually labelled particle image data of a diverse list of proteins is a key bottleneck hindering the development of machine learning and AI methods to automate protein particle picking.

Creating a high-quality manually labelled single-protein particle dataset of a large, diverse set of representative proteins to facilitate machine learning is a challenging task. Single-particle cryo-EM images suffer from high background noise and low contrast due to the limited electron dose to minimize the radiation damage to the biomolecules of interest during imaging, which makes particle picking difficult even for human. Low signal-to-noise ratio (SNR) of the micrographs, presence of contaminants, contrast differences owing to varying ice thickness, background noise fluctuation, and lack of well-segregated particles further increases the difficulty in particle identification ^30^. This is one reason there is still a lack of large manually curated protein particle datasets in the field.

A common problem of the particle picking algorithms trained on a small amount of particle data of a few proteins is that they cannot distinguish ‘good’ and ‘bad’ particles well, including overlapped particles, local aggregates, ice contamination and carbon-rich areas ^31^. For instance, the methods: DRPnet ^32^, TransPicker ^33^, CASSPER ^34^, and McSweeney et al.’s method ^35^ that made significant contributions to the particle selection problem suffered the two similar problems. Firstly, there is not a sufficient and diversified dataset to train them. Secondly, there is no gold standard to test them. The similar problems happened to other supervised and unsupervised machine learning methods, such as an unsupervised clustering approach ^36^, AutoCryoPicker ^27^, DeepCryoPicker ^28^, APPLE picker ^18^, Mallick et al’s method ^37^, gEMpicker ^38^, Langlois et al.’s method ^31^, DeepPicker ^19^, DeepEM ^20^, Xiao et al.’s method ^21^, APPLE picker ^18^, Warp ^25^, SPHIRE-crYOLO ^39^, and HydraPicker ^40^ all encountered similar problems. They usually perform well on the small, standard datasets used to train and test them (e.g., Apoferritin and KLH), but may not generalize well to non-ideal, realistic datasets containing protein particles of irregular and complex shapes, which are generated daily by the cryo-EM facility around the world.

To address this key bottleneck hindering the development of machine learning and AI methods for automated cryo-EM protein particle picking, we created a large dataset (CryoPPP) of cryo-EM micrographs in which protein particles were manually labelled. The micrographs are associated with 32 representative proteins of diverse sequences and structures that cover a much larger protein particle space than the existing datasets of a few proteins such as Apoferritin and KLH. The quality of the manually labeled particles of selected proteins was rigorously validated against some particles labelled by the authors who generated the cryo-EM data by both 2D particle class validation and 3D cryo-EM density map validation. The quality of our manual annotation is in pair with the annotations provided by the experts who created the data in the first place, which confirms our manual particle labelling process is effective. Therefore, we believe CryoPPP is a valuable resource for training and testing machine learning and AI methods for automated protein particle picking.

## Methods

CryoPPP was created through a series of steps as shown in **Figure 2**. We first crawled the data from the EMPIAR website using API and FTP scripts. We filtered out microscopic images of various non-single-protein particles (e.g., bacteria, filaments, RNA, protein fibril, virus-like particles) and retained only high-resolution micrographs acquired by cryo-EM technique for manual particle labeling.

**Figure 2:**
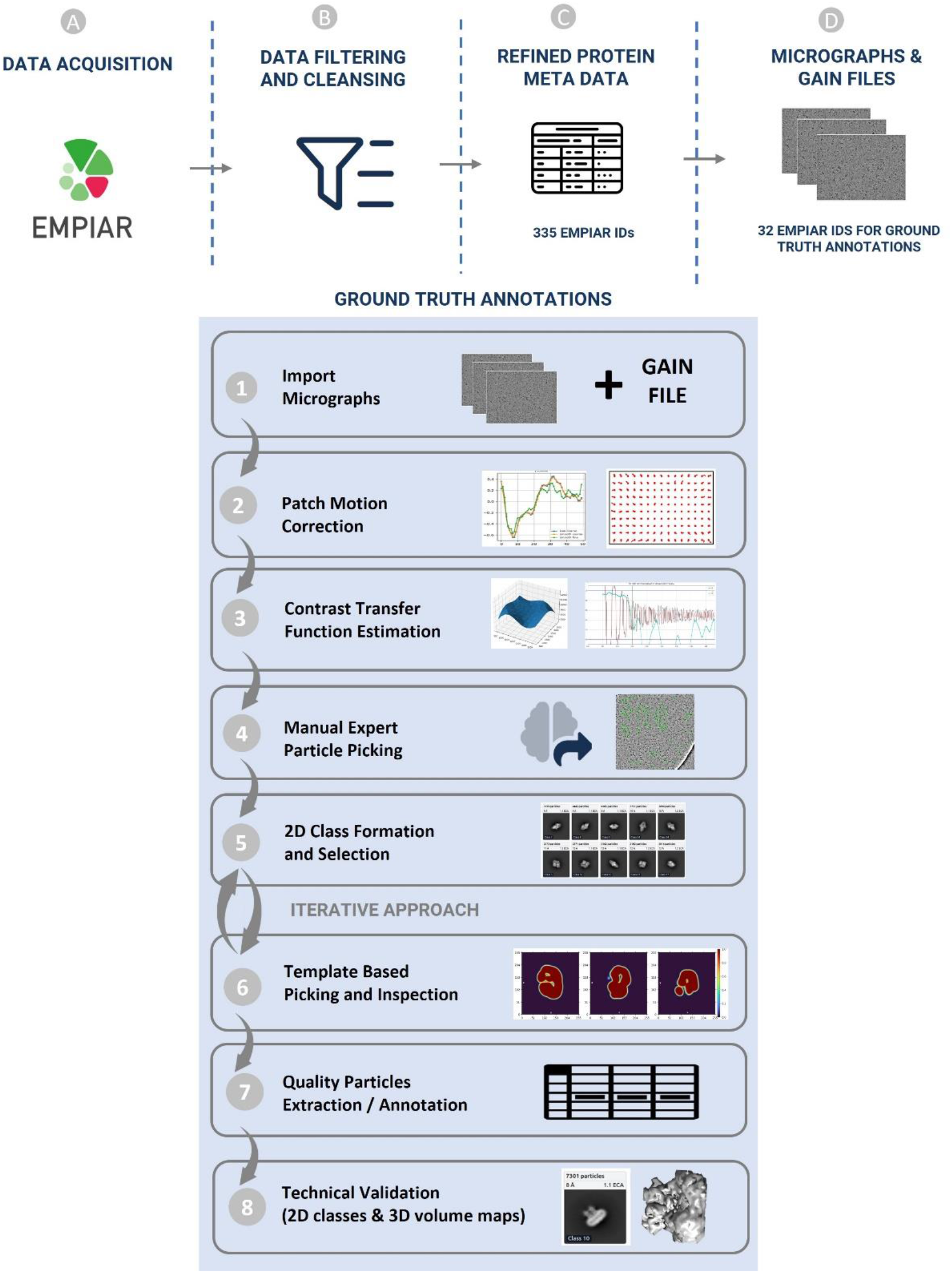
Graphical illustration of the overall methodology of creating CryoPPP dataset with manual EMPannotations of protein particles. (A)-(D) represent the steps for data acquisition and protein metadata preparation. (1)-(8) represent subsequent steps for the ground truth annotation and validation of picked protein particles. The iterative approach between step (5) and (6) is carried out to achieve the high-quality picking of particles.

After importing the micrographs with all the physiochemical parameters gathered from the corresponding published literature, we performed motion correction and Contrast Transfer Function (CTF) estimation for them. Once the micrographs were prepared, two human experts manually picked the particles after setting up the low pass filter values and proper diameter for picking particles.

The expert-picked particles were cross-validated and then went through 2D particle classification. The best particles based on resolution, particle count, and visually appealing and sensible 2D classes were selected and further used for template-based particle picking and further human inspection. After iterating the 2D classes from template-based picking and human inspection, we ultimately obtained the final set of highly confident protein particles as ground truth and exported them in the files in star, csv and mrc formats. The first two files (.star and .csv) contain the coordinates of the protein particles and the latter (.mrc) store particles stacks. The process of creating CryoPPP in **Figure 2** is described in the following sections in detail.

### I EMPIAR Metadata Collection and Filtering

The process of preparing the dataset began with collecting metadata about cryo-EM image datasets in EMPIAR. Data collection scripts that use API and FTP protocols were used to automatically download the metadata from the EMPIAR web portal ^29^. The metadata includes EMPIAR ID of each cryo-EM dataset of a protein, the corresponding Electron Microscopy Data Bank (EMDB) ID, Protein Data Bank (PDB) ID, size of dataset, resolution, total number of micrographs, image size/type, pixel spacing, micrograph file extension, gain/motion correction file extension, FTP path for micrograph/gain files, Globus path for micrograph/gain files, and publication information.

Following the metadata collection, the individual cryo-EM datasets in the collection were filtered as depicted in **Figure 3 (Steps 1 - 5)**. First, we only chose EMPIAR IDs (datasets) that have their volume maps deposited in EMDB. From the chosen EMPIAR datasets, we only selected ones that had corresponding protein structures in the Protein Data Bank (PDB).

**Figure 3:**
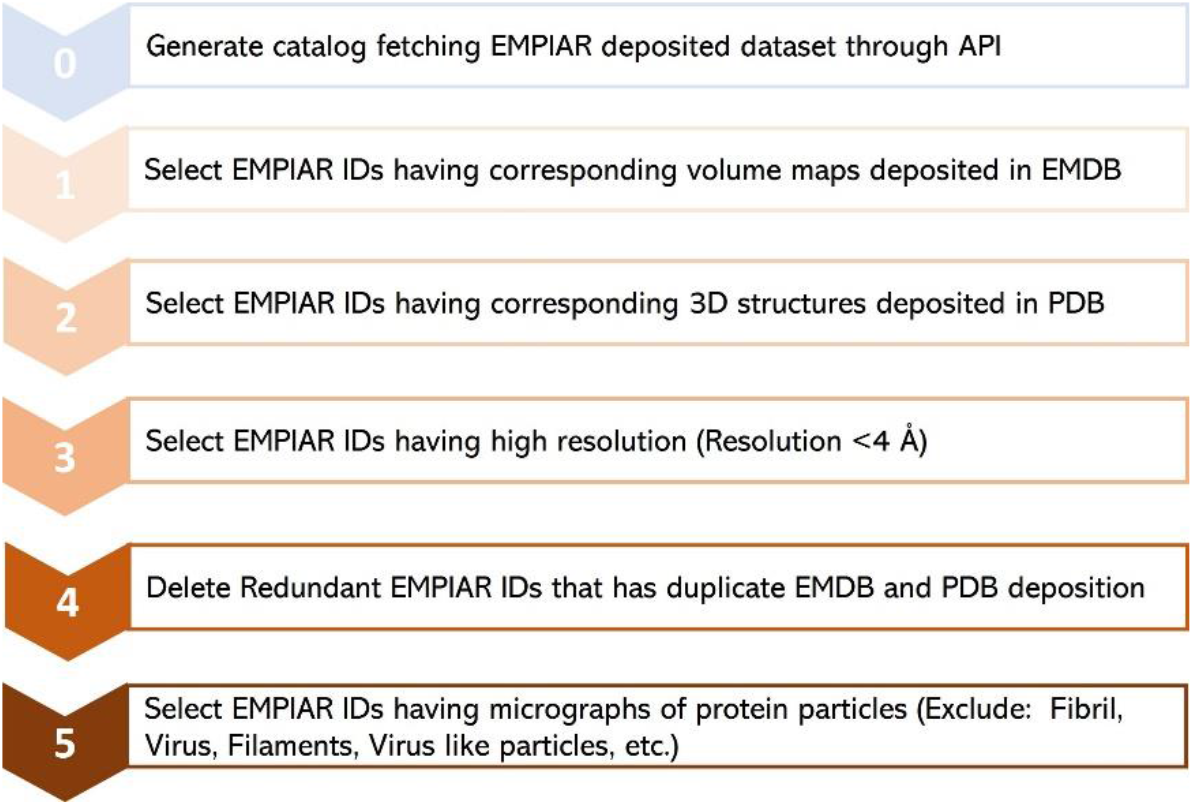
The step-by-step procedure for collecting and selecting Cryo-EM protein datasets from EMPIAR database. 335 unique EMPIAR datasets (IDs) of 335 proteins were selected at the end.

To ensure high data quality, we then retained only the EMPIAR datasets whose resolution was less than 4 Angstrom (Å). We observed that there were some redundant EMPIAR datasets (e.g., EMPIAR ID: 10709 & 10707, EMPIAR ID: 10899 & 10897) that correspond to the same biomolecule with the same PDB and EMDB IDs. Hence, we eliminated those duplicate entries. After removing duplicate records, we selected only EMPIAR datasets that contained micrographs of protein particles, excluding other biological samples such as viruses. This filtering step required some literature study of individual EMPIAR datasets. The motion correction and gain correction files for the selected datasets were extracted from the EMPIAR if required. The final list of meta data includes 335 EMPIAR entries, 32 out of which were used for manual labelling. Refer to the *EMPIAR_metadata_335*.*xlsx* file in CryoPPP for further information about the list of 335 datasets of 355 proteins.

### II Manual Particle Labeling

Manually picking particles in cryo-EM micrographs through the GUI interfaces of cryo-EM analysis tools such as CryoSPARC ^16^, EMAN2 ^14^ and RELION ^15^ is very time consuming. Additionally, it is highly challenging to import micrographs, carry out motion correction, and estimate CTF for large micrographs. Furthermore, it takes a lot of disk space to store the labelled particle data together with the corresponding micrographs and particles stack files. Therefore, we chose 32 representative EMPIAR datasets out of 335 entries selected in the previous section for manual particle labelling to create the CryoPPP dataset, considering diverse particle size/shapes, density distribution, noise level, and ice and carbon areas. Moreover, proteins from a wide range of categories, such as: metal binding, transport, membrane, nuclear, signaling, and viral proteins were selected. See supplementary Tables 1 and 2 for more details about the 32 proteins (cryo-EM datasets). Most of the pre-processing, manual particle labelling, real-time quality assessment, and decision-making workflows were performed using CryoSPARC v4.1.1 ^16^, EMAN2 ^14^, and RELION 4.0 ^15^.

CryoPPP includes a total of 9,089 micrographs (∼300 Cryo-EM images per selected EMPIAR dataset). We labelled ∼300 micrographs per EMPIAR data because using all the micrographs in each dataset would result in 32.993 TB of data, which would be too big for most machine learning tasks. Another reason is that many micrographs in the same EMPIAR dataset are similar and therefore it is not necessary to include all of them. The particle labelling process is described in detail as follows.

#### 1 Importing Movies

This is the crucial first step of particle labeling. For each EMPIAR dataset, we import two inputs: micrographs and gain reference (motion correction files). We analyzed the description of the EM data acquisition and grid preparation for each dataset in order to collect the important information such as raw pixel size (Å), acceleration voltage (kV), spherical aberration (mm), and total exposure dose (e/Å ^2^) for the micrographs in the dataset.

Furthermore, we obtained gain reference for micrographs if their motion was not corrected before. We used *e2proc2d*, a generic 2-D image processing program in EMAN2 ^14^, to convert different formats of motion correction file (e.g., .dm4, .tiff, .dat, etc.) to .mrc file since CryoSPARC accepts only .mrc extension. Then, based on the microscope camera settings and how the data was acquired during the imaging process, we applied geometrical transformations (flip gain reference and defect file left-to-right/top-to-bottom (in x/y axis) or rotate gain reference clockwise/anti-clockwise by certain degrees) relative to the image data. **Supplementary Table S1** contains the details of input parameters for each EMPIAR ID. After importing movies and motion correction files, we proceeded to the job inspection panel of CryoSPARC to ensure that all input settings and loaded micrographs were correct.

#### 2 Patch Motion Correction

When specimens are exposed to an electron beam, the mobility of sample molecules (protein particles) during data acquisition can affect the overall quality of electron micrographs and lower the final resolution ^41^. Hence, it is necessary to correct the movement of particles (referred to as ‘beam-induced motion’).

The causes behind this motion can be categorized into two types: (1) *Motion from Microscope:* It is caused by stage drift and usually occurs in microscope due to little amount of vibration left over after the stage has been aligned to a new position ^42^. It moves the sample relative to the beam and optical axis. This motion is quite jagged in time, with sharp accelerations or twitches, but is consistent. The entire image will move in the same direction over time. (2) *Motion from sample deformation:* This motion is caused by the energy deposited into the ice by the beam, or energy already trapped in it, due to strained forces locked in during freezing. It is eventually released during the image capturing process. As the electrons pass through the samples, the energy from the beam and the temperature change causes the ice to physically deform and bend. That deformation is often smoother over time, but it can be highly anisotropic in space. In this case, various parts of the same image can move in different directions at the same time.

Both motions must be estimated and corrected to obtain high-resolution reconstructions from the data. In the patch-based motion correction step, we corrected both global motion (stage drift) and local motion (beam-induced anisotropic sample deformation) for the micrographs (as shown in **Figure 4 (A)**. In the anisotropic deformation plot in **Figure 4 (B)**, each red circle indicates the center of a single “patch” of the image, and the curves emerging from each circle show the motion of that portion of the sample. We can observe the correlation between the motion of adjacent patches. They move somewhat similarly to one another. To prevent the fit from being distorted by random noise in the micrograph, the patch motion correction algorithm imposes smoothness constraints on the motion.

**Figure 4:**
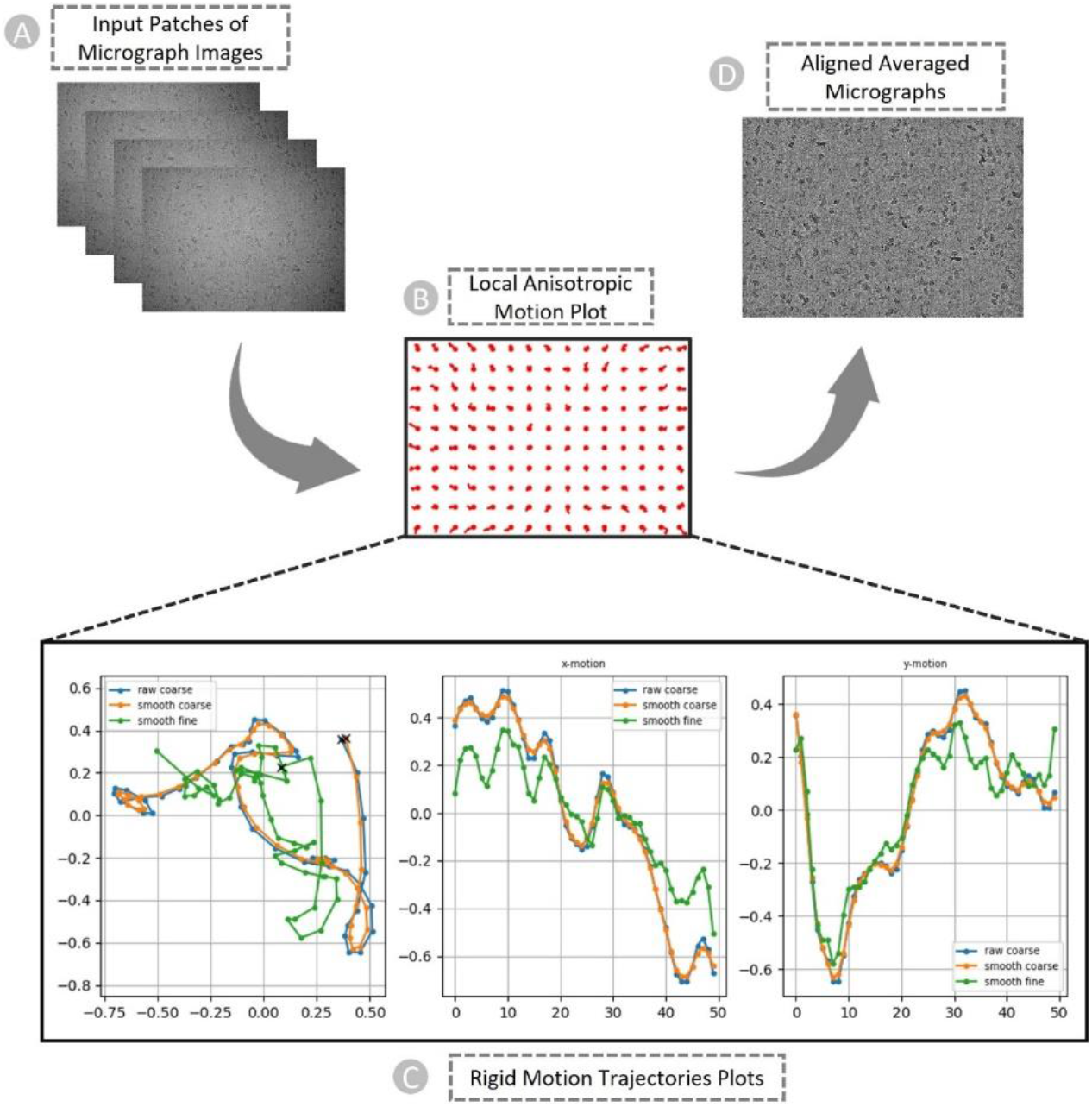
The patch-based local and global motion correction pipeline for EMPIAR ID 10737 (E. coli cytochrome bo3 in MSP Nanodiscs). (A) Full frames of micrographs as input. (B) Anisotropic deformation. (C) Rigid motion trajectories plots. Blue: original trajectory, Radish: trajectory with small smoothing penalty, Green: trajectory with fine smoothing. Left: Overall motion trajectory over X and Y motion. Center: X-motion plot over time. Right: Y-motion plot over time. (D) Non-dose weighted aligned averaged micrographs with the highest amount of signal and least amount of motion blur as output.

**Figure 4 (C)** are the examples of plots generated by patch motion correction that depict the computed trajectories. The set of plots shows overall motion correction (an actual trajectory plot, followed by X-motion plot and Y-motion plots over time). In the overall motion trajectory over X and Y motion (**Figure 4 (C), Left**), each dot represents the sample’s position from frame to frame. Here, the x and y axes represent the units of pixels in the raw data’s pixel size. The sample begins at point (**X**), moves downward, makes a curve and again changes direction toward the left-top, and then continues to descend to the left. We apply this trajectory to the input data by shifting each image in reverse of what the motion trajectory suggests and finally averaging images together. In other words, we track a sample’s motion during the exposure to undo it.

#### 3 Patch-based CTF Estimation

The contrast of images captured in the electron microscope is affected by imaging defocus and lens aberrations, which are adjusted by microscope operators to enhance the contrast. The relationship between lens aberrations and the contrast in the image is defined by the CTF. It explains how information is transferred as a function of spatial frequency.

It is important to estimate CTF, which is then corrected during 2D particle classification and 3D reconstruction steps. Otherwise, the feasible reconstruction will have extremely low resolution. A full treatment of the effects of the CTF usually proceeds in two stages: CTF estimation and CTF correction. In CryoSPARC the CTF model is given by the equation I.

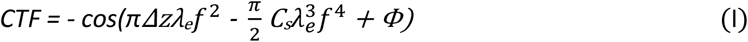

where *ΔΖ*is defocus, *λ*_*e*_ is the wavelength of the incident electrons, *C*_*s*_ *is* spherical aberration, and *f* is spatial frequency. *Φ*represents a phase shift factor.

Most cryo-EM samples are not ‘flat’. Before a sample is frozen, particles tend to concentrate around the air-water interfaces, and the ice surface itself is usually not flat ^43,44^. Because defocus has an impact on the CTF, distinct particles can have various defoci and hence various CTFs within a single image. To address this problem, CryoSPARC offers a patch-based CTF estimator that analyzes numerous regions of a micrograph to calculate a “defocus landscape”.

We performed a 1D search over defocus for every micrograph. **Figure 5 (A)** depicts the 1D search for a particular micrograph of EMPIAR ID 10737 ^45^. This plot helps identify a particular defocus value that stands out among a variety of other defocus values (x-axis). Patch CTF creates a plot showing how closely the input micrographs’ observed power spectrum and the calculated CTF match. The CTF fit plot in **Figure 5 (B)** shows that the computed CTF matches the observed power spectrum up to a resolution of 3. 993 Å. The cross correlation between the observed spectrum and the calculated CTF is depicted by the cyan line in the plot. The vertical green line in the plot represents the frequency at which the fit deviates from CryoSPARC’s cross-correlation threshold of 0.3 for a successful fit.

**Figure 5:**
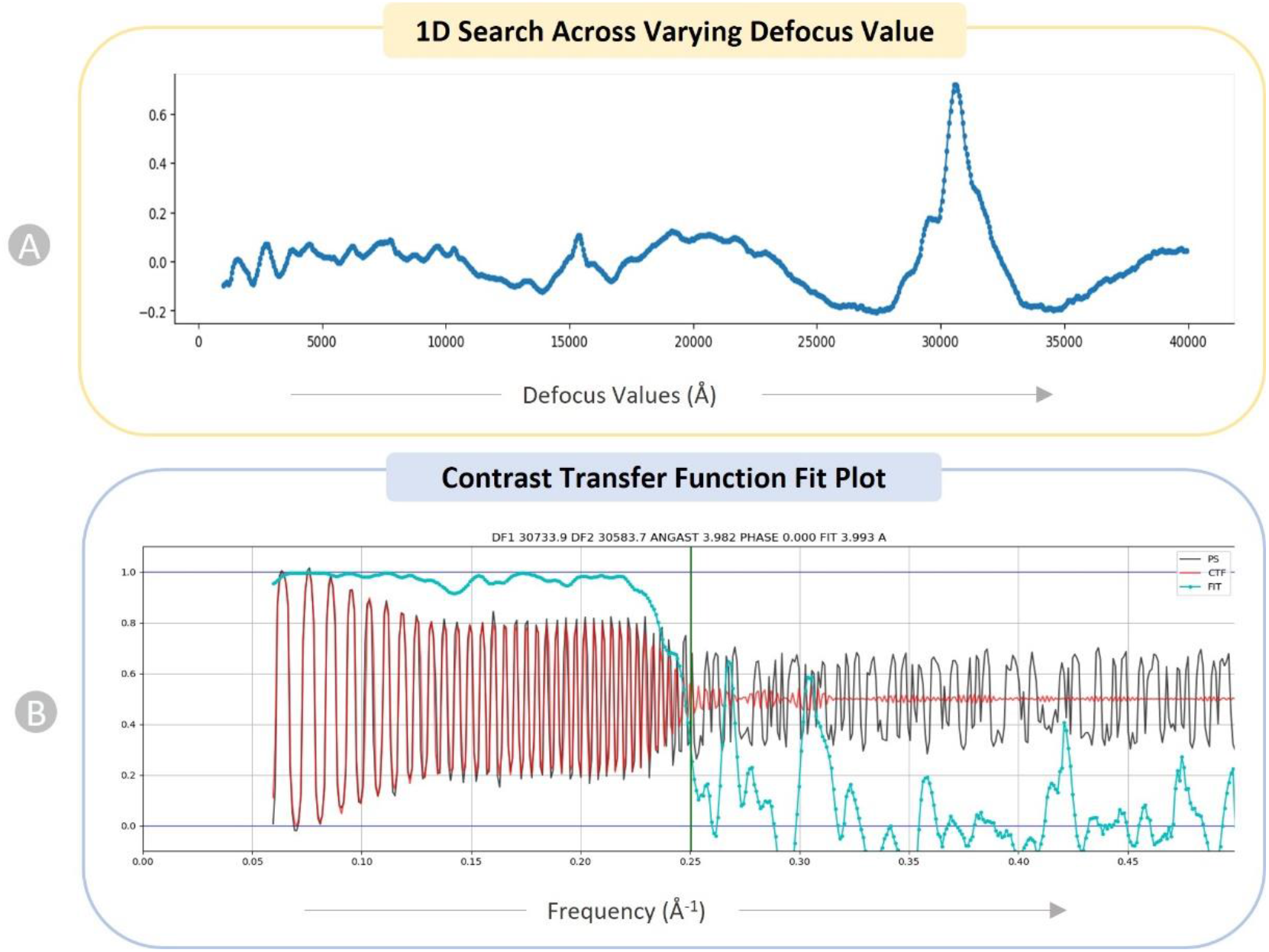
Diagnostic plots of CTF for EMPAIR 10737 (E. coli cytochrome bo3 in MSP Nanodiscs). (A) 1D search over varying defocus values (underfocus). (B) CTF fit plot. X-axis displays frequency, in units in inverse angstroms (Å^-1^) and Y-axis shows correlation metric between power spectrum (PS) and CTF value. Black: observed experimental power spectrum. Red: calculated CTF. Cyan: cross-correlation (fit).

We executed the patch CTF to obtain the output micrographs with data on their average defocus and the defocus landscape. When particles were extracted, this data was automatically used to assign each particle a local defocus value based on its position in the landscape.

#### 4 Manual Particle Picking

After performing the motion correction and CTF estimation, we manually picked particles interactively from aligned/motion-corrected micrographs with the goal of creating particle templates for auto-picking. Depending on the size and shape of the protein particles, we adjusted the box size and the particle diameter. Since picking particles on raw micrographs is extremely difficult, we tweaked the ‘Contrast Intensity Override’ while viewing micrographs in order to obtain the best distinctive view for picking particles.

It is particularly challenging to manually pick particles from micrographs with smaller defocus levels, and vice versa. **Figure 6** illustrates the visualization of micrographs in the same dataset with different defocus levels for EMPAIR 10532 ^45^. Hence, to generate comprehensive templates from a wide range of defocus values, we manually picked particles from multiple micrographs with diverse defocus and CTF fit values.

**Figure 6:**
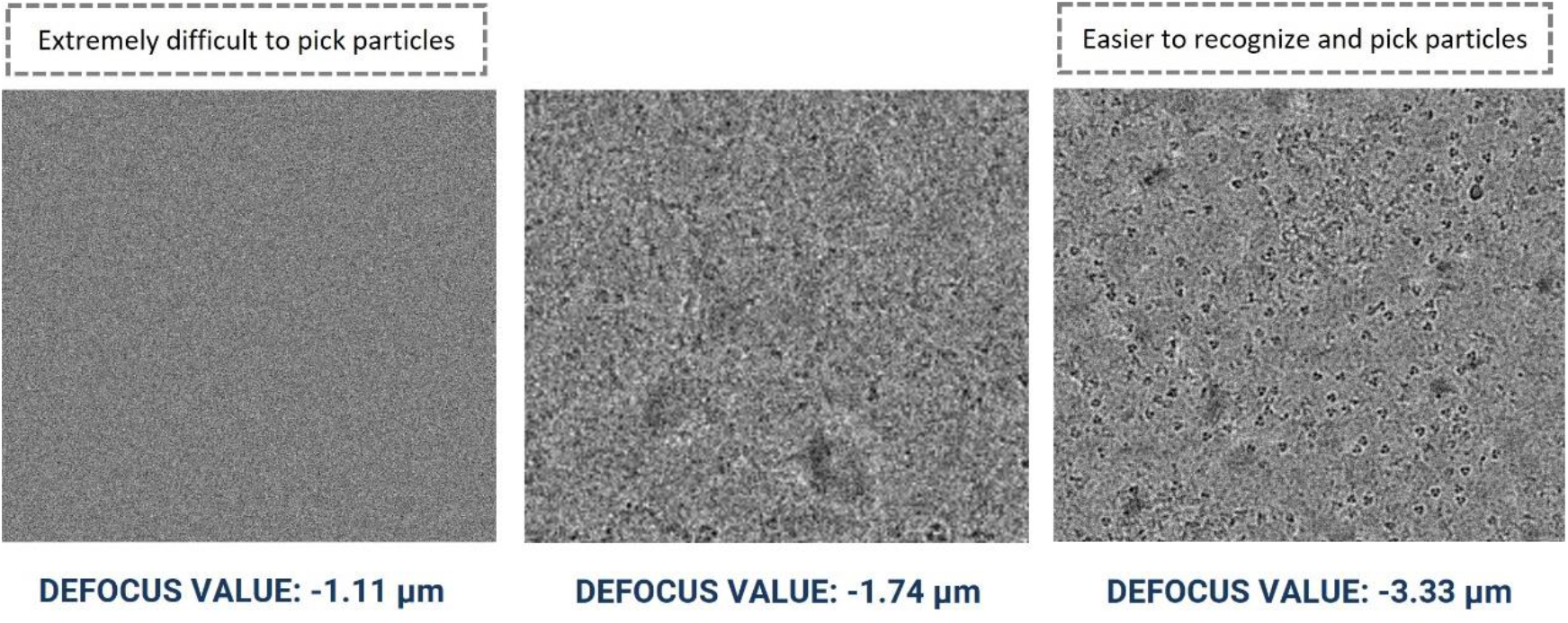
Cryo-EM micrograph images of EMPIAR ID 10532 (Influenza Hemagglutinin) with different defocus values. Micrographs with smaller defocus values make particle picking difficult and vice-versa.

As manual picking was very time intensive, we selected a subset of micrographs (around 20 micrographs of each EMPIAR dataset) for manually picking initial particles for the subsequent template-based particle picking. More details regarding the total number of particles picked manually including the total number of micrographs considered for manual pick are provided in the **Supplementary Table S2**.

#### 5 Forming and Selecting Best 2D Particle Classes

The manually picked particles went through the 2D classification step. This step helped to classify the picked particles into several 2D classes to facilitate stack cleaning and junk particles removal. To analyze the distribution of views within the dataset qualitatively, we specified a specific number of 2D classes. By doing this, we investigated the particle quality and removed junk particle classes, which ultimately facilitated the selection of good particle classes.

We specified the initial Classification Uncertainty Factor (ICUF) and maximum alignment resolution to align particles to the classes with 40 expectation maximization (EM) iterations. The diameter of the circular mask that was applied to the 2D classes at each iteration was controlled using the circular mask diameter in the case of crowded particles.

After the 2D classes were formed, we selected the best particle classes interactively to remove the junks. **Figure 7** shows an example of 2D classification and selection of highly confident particles for EMPIAR ID 10017 ^46^. We used three diagnostic measures to select the 2D classes: resolution (Å) of a class, the number of particles of a class (higher, better), visual appearance of a class. Considering only the number of particles in a class is not sufficient because some classes containing a small number of particles may represent a unique view of the protein.

**Figure 7:**
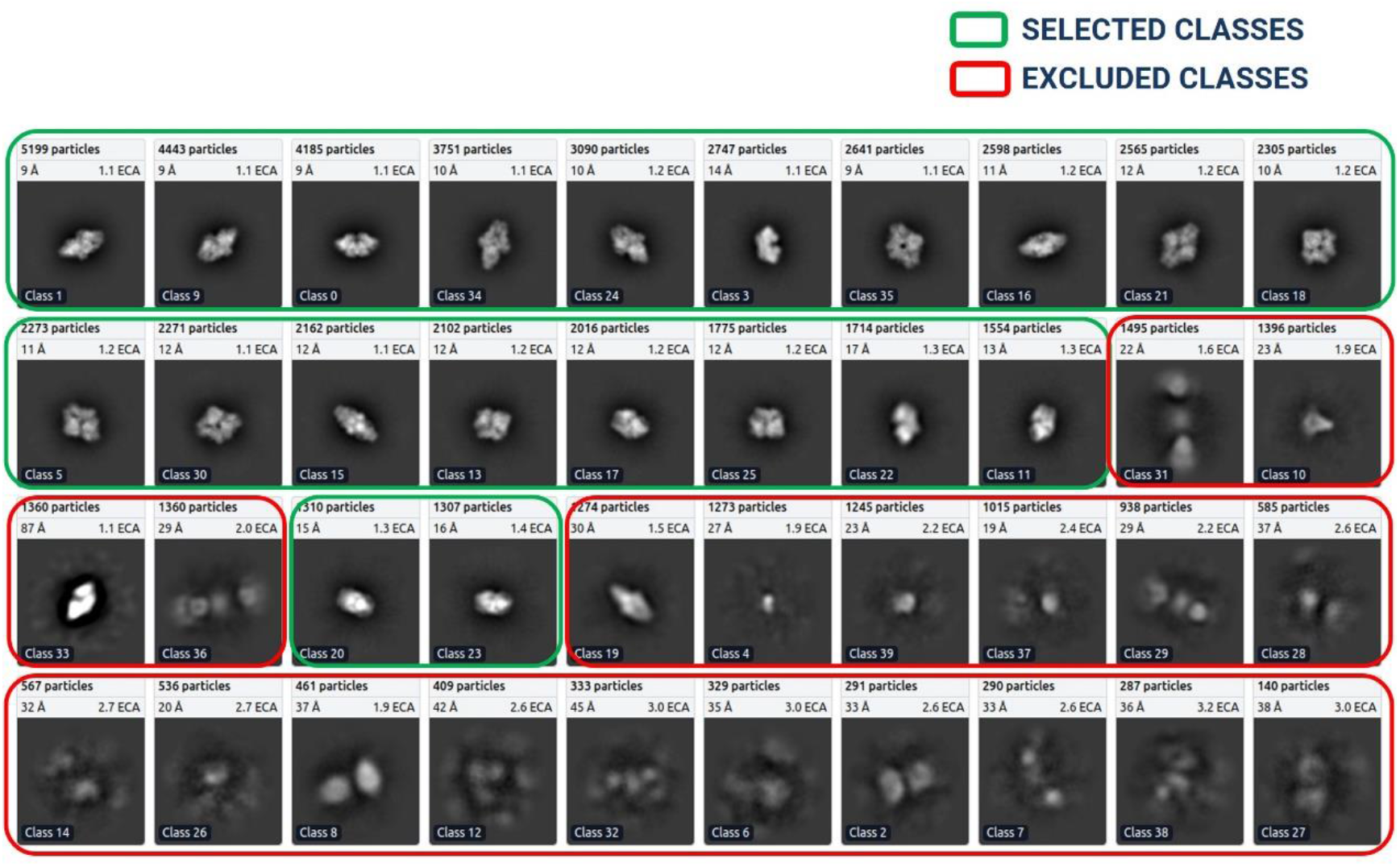
2D classes for EMPIAR ID 10017 (Beta-galactosidase), ordered ascendingly by the number of particles assigned to each class. Green: High quality particle classes selected for further template-based picking. Red: Rejected particle classes.

#### 6 Template based Picking and Manual Inspection and Extraction of Particles

After the best particle classes were selected and exported, we used a template generated from the ‘**Forming and Selecting Best 2D** ‘ step to pick more particles. The process was iterative, meaning that the output of a round of ‘template-based picking and inspection’ was again utilized for ‘2D class formation’ step to form and select best 2D classes under the human inspection. This process was repeated until we acquired high resolution particles that include all possible particle projection angles.

The final templates with green boxes (as shown in **Figure 7**) were used to execute auto-pick particles from micrographs. With CryoSPARC’s Template Picker, we used high resolution templates to precisely select particles that matched the geometry of the target structure. **Figure 8 (A)** represents manually picked particles for EMPIAR-10017 ^46^ that work as templates to facilitate template-based picking that eventually results in template-based picked particles ready for human inspection as shown in **Figure 8 (B)**. We specified constraints like particle diameter in angstrom (see **Supplementary Table S2** for more information) and a minimum distance between particles to generate the templates based on the SK97 sampling algorithm ^32^ to remove any signals from the corners and prevent crowding. We observed that the blob-based in picking in RELION required minimum and maximum allowed diameter of the blobs, whereas defining a single value for particle’s diameter worked well in CryoSPARC.

**Figure 8:**
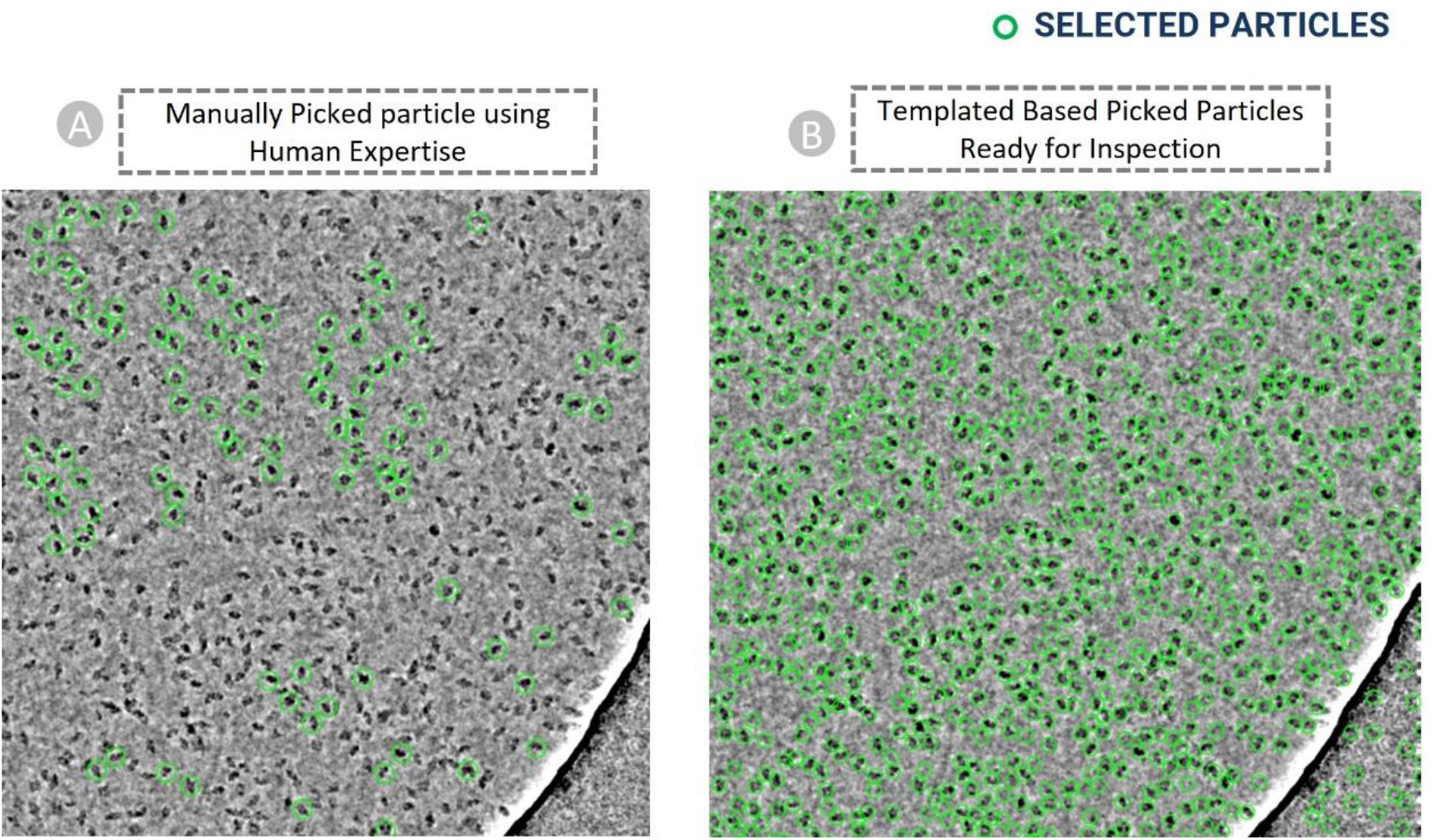
Cryo-EM micrograph image of EMPIAR ID 10017 (defocus value: -3.63 µm) used for template-based particle picking. (A) Micrograph with manually picked protein particles (encircled with green circle, particle diameter: 190 Angstrom, low pass filter value: 25). (B) Picked protein particles with template-based picking ready for manual inspection and the adjustment of power value and NCC score.

Finally, the particles obtained by the template picking went through the manual inspection step, where we examined and modified picks using various thresholds. We adjusted the lowpass filter, normalized cross-correlation (NCC), and power threshold to improve the visibility of the picks and removed false positives as shown in **Figure 11 (B)**. The 2D colored histogram plots as depicted in **Figure 11** were used to scrutinize micrograph median pick scores versus defocus for extracting the coordinates of high-quality protein particles.

## Data Records

The CryoPPP dataset consists of manually labelled 9,089 micrographs of 32 diverse, representative cryo-EM datasets of 32 protein complexes selected from EMPIAR. Each EMPIAR dataset identified by a unique EMPIAR ID has about ∼300 cryo-EM images in which the coordinates of protein particles were labeled and cross-validated by two experts aided by software tools.

Each data folder (named by its corresponding EMPIAR ID) includes the following information: original micrographs (either motion-corrected or not), gain motion correction file, new motion-corrected micrographs (if original micrographs are not motion-corrected), ground truth labels (manually picked particles), and particles stack. The directory structure of each data entry is illustrated in **Figure 9**. The data in each directory is described as follows. It is worth noting that if the original micrographs were not motion-corrected, we applied the motion correction to them to create their motion-corrected counterparts.

**Figure 9:**
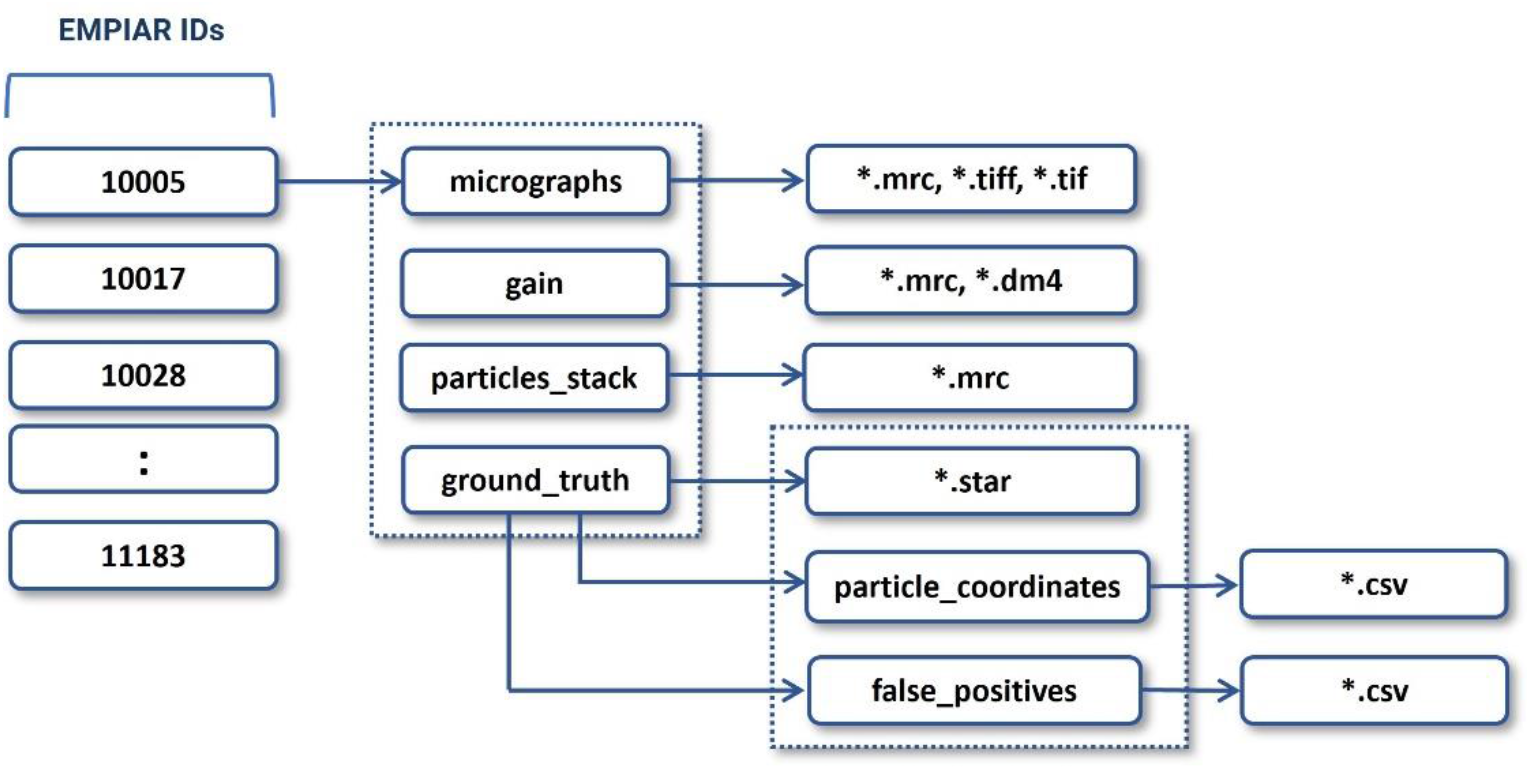
The directory structure of each expert-labelled data entry of CryoPPP. The directory contains micrographs, motion correction files, particle stacks, and ground truth labels (manually picked particles). The blocks with numbers on the left represent corresponding EMPIAR IDs.

### I. Raw Micrographs

These are the two-dimensional projections of the protein particles in different orientations stored in different image formats (MRC, TIFF, EER, TIF, etc.). They can be considered as the photos taken by cryo-EM microscope. Original micrographs are from EMPIAR and can be either motion corrected or not. If an entry has a ‘*gain’* folder, it includes both raw non-motion-corrected micrographs and their motion-corrected counterparts created by us. Users are supposed to use the motion corrected micrographs as input for machine learning tasks. The scripts for the motion correction are available at CryoPPP’s GitHub website.

### II. Motion Correction (gain files)

It contains motion correction files (if motion in original micrographs not corrected before) stored in different formats like dm4 and mrc. It is used to correct both global motion (stage drift) and local motion (beam-induced anisotropic sample deformation) that occur when specimens (protein particles) are exposed to the electron beam during imaging. Correcting the motion enables the high-resolution reconstruction from the data.

### III. Particles Stack

Particle stack comprises of the mrc files (with names corresponding to individual micrographs’ filenames) of manually picked protein particles (ground truth labels). These are three-dimensional grids of voxels with values corresponding to electron density (i.e., a stack of 2D images). To browse and examine this file, utilize EMAN2 ^14^, UCSF Chimera ^47^, or UCSF ChimeraX ^48^.

### IV. Ground Truth Labels

Ground truth data contain the star and CSV files for both all true particles (positives) and some typical false positives (e.g., ice contaminations, aggregates, and carbon edges). The positive star (and corresponding CSV) files are the ground truth position of the picked particles combined in a single file for all ∼300 micrographs per EMPIAR ID. While the negative star file consists position of the false positive particles. These star files contain information like X-coordinate, Y-coordinate, Angle-Psi, Origin X (Ang), Origin Y (Ang), Defocus U, Defocus V, Defocus Angle, Phase Shift, CTF B Factor, Optics Group, and Class Number of the particles.

Besides, there is a subdirectory called *particle_coordinates* inside *ground_truth*, which contains csv files, with same name as raw micrographs, which contain individual protein particle’s X-Coordinate, Y-Coordinate along with their diameter and other relevant information.

## Technical Validation

To ensure that the dataset is of high quality, we applied numerous validations and statistical analyses throughout the data curation process.

### I. Quality of Data

As noted in **Figure 3**, we ensure that the dataset exclusively contains micrographs obtained using the Cryo-EM technique. Only the EMPIAR IDs with resolution less than 4 Å are chosen for creating refined protein metadata and ground truth labels of protein particles. The detailed quality control procedures are described as follows.

### II. Distribution of Data

#### a) Diverse Protein Types

To be inclusive and ensure unbiased data generation, we selected the cryo-EM data of 32 different, diverse protein types (e.g., membrane, transport, metal binding, signalling, nuclear, viral proteins) to manually label protein particles, which can enable machine learning methods trained on them to work for many different proteins in the real-world. We selected the datasets covering different particle size, distribution density, noise level, ice and carbon areas, and particle shape as they are influential in particle picking.

#### b) Diverse Micrographs within the Same Protein Type

The variance in micrographs’ defocus values within a EMPIAR dataset is not accounted for by majority of the particle picking methods. This defocus variation causes the same particles to appear differently, altering the noise statistics of each micrograph. This makes it challenging to create thresholds to select high quality particles. **Figure 6** shows an example how different defocus values impact the appearance and quality of Cryo-EM images in the same EMPIAR dataset. Therefore, during manually picking the particles, we included a wide variety of defocus levels and CTF fit.

We recorded the correlation between defocus levels and the pick scores / the power scores (shown in **Figure 10** for EMPIAR**-**10590 ^48^) to assess the shape and density of a particle candidate independently. After calibration, the scores of each particle are recorded relative to the calibration line, and these values are used to define thresholds on the parameters.

**Figure 10:**
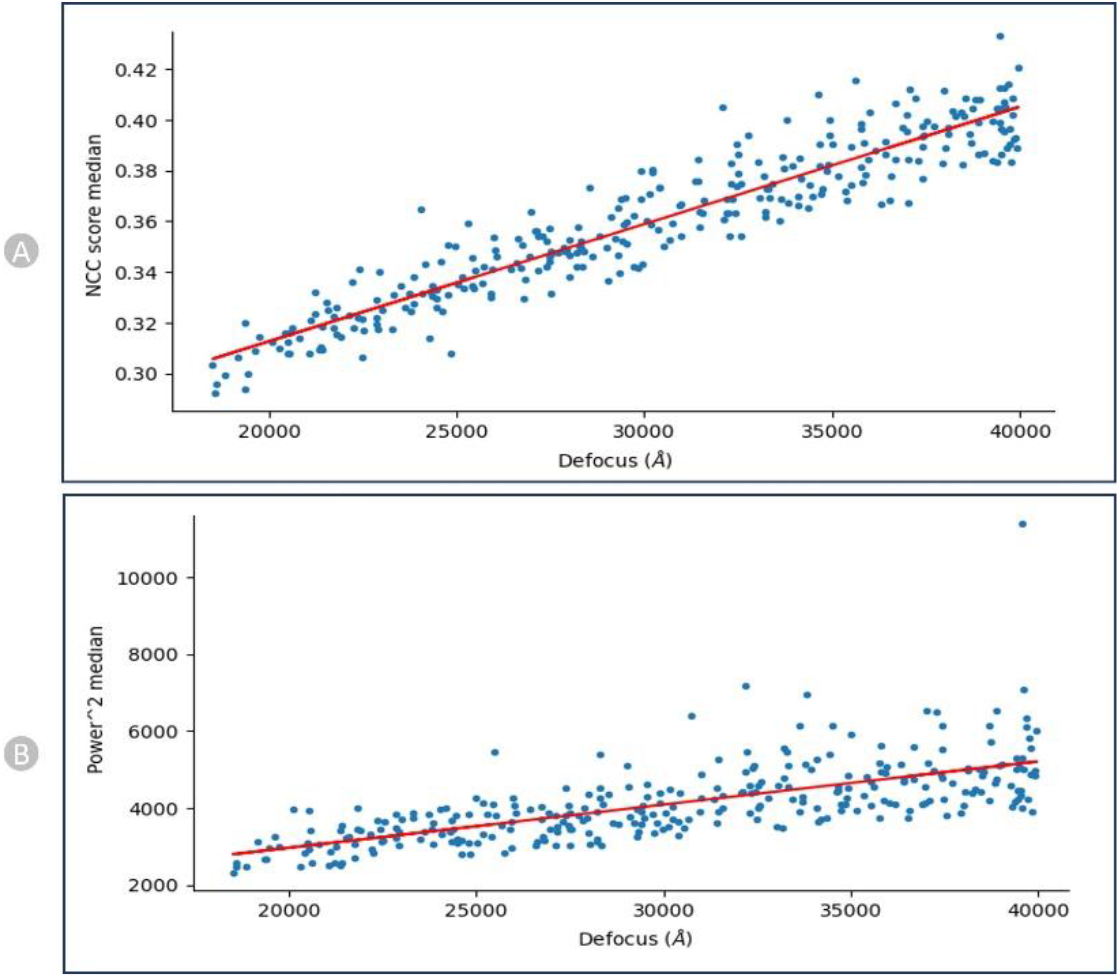
NCC and Power calibration plots for EMPIAR-10590 (Endogenous Human BAF Complex). (A) Calibrating Median NCC scores vs defocus. (B) Calibrating Power scores vs Defocus. There is a strong trend that higher defocus correlates with higher NCC scores and same with Power score.

### III. Reliability of Ground Truth Annotations

#### a) Legitimacy of Importing Micrographs and Motion Correction Data

All the input parameters used to prepare for loading micrographs into the CryoSPARC system were gathered from the appropriate literature. We adhered to the standards in the publications including data acquisition and imaging settings such as the microscope used, defocus range, spherical aberration, pixel spacing, acceleration voltage, electron dose and the correct usage of motion correction. Based on the microscope settings during the imaging process, we applied appropriate geometrical transformations. The defect files and the motion-correction files were flipped left-to-right or top-to-bottom and also rotated by specific degrees in clockwise/anti-clockwise direction as required. All these factors were thoroughly investigated and used during the data loading process in CryoSPARC.

#### b) Inspection of Picked Protein Particles

The picked particles were inspected using a 2D colored histogram, as shown in **Figure 11**. A particle of interest would have an intermediate local power score and a high template correlation (indicating its shape closely matches its template). Low local power scores indicate empty ice patches, even though it might resemble the template. Additionally, very high local power scores indicate carbon edges, aggregates, contaminants, and other objects with excessive densities that resemble particles.

**Figure 11:**
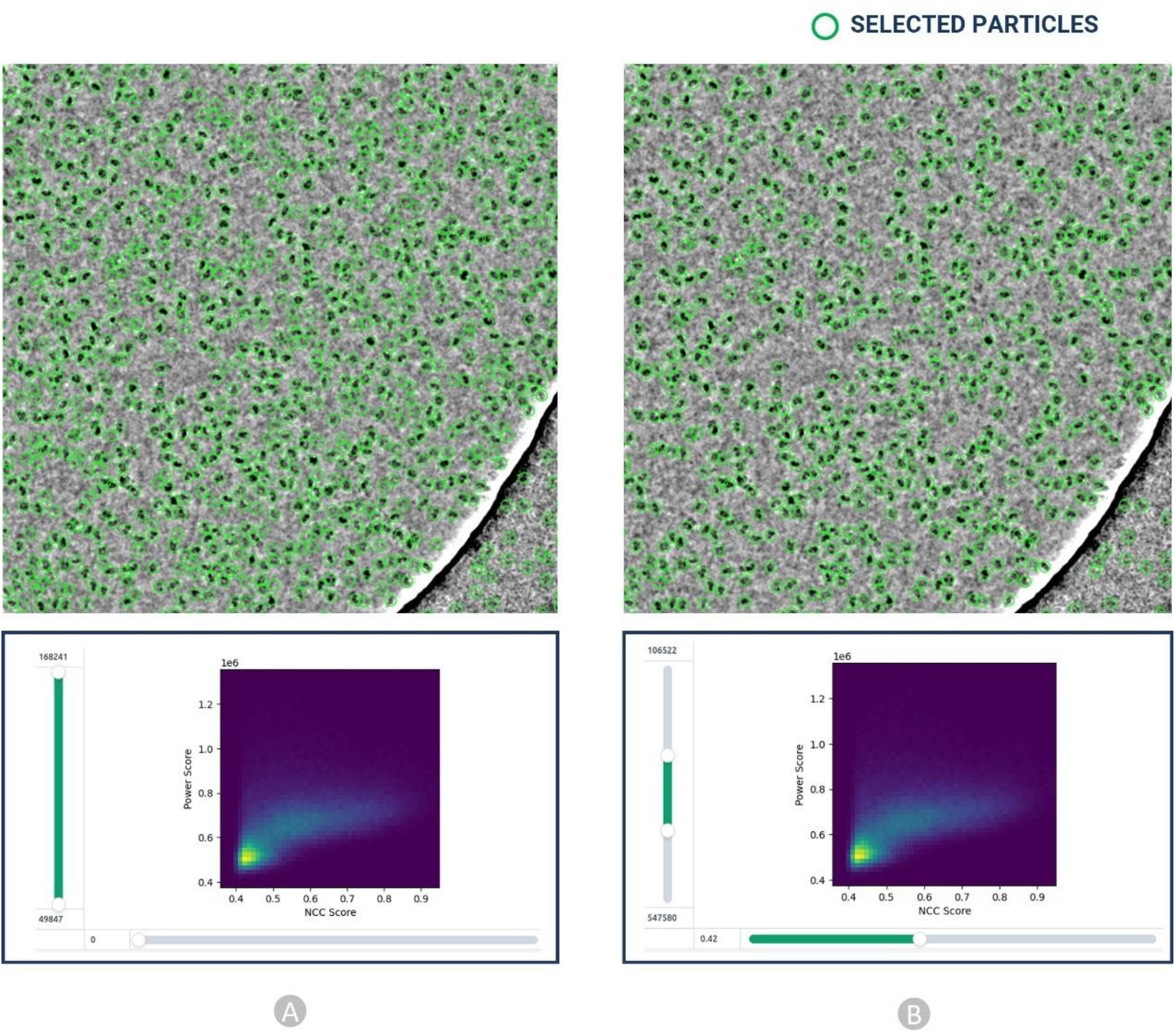
Particle inspection and filtration by adjusting normalized cross correlation (NCC) score (X axis) and local power (Y axis) for EMPIAR 10017. (A) Initial picked particles (green circles) from template-based picking step. (B) Selected high quality protein particles through adjustment of NCC and power score values.

As shown in **Figure 11 (B, bottom)**, we interactively specified the upper and lower thresholds for both the Power score and NCC score for each dataset improving the accuracy in the manual particle picking.

#### c) Cross-validation by two Human Experts

The results of the particles picked by the two Cryo-EM experts were compared to each other to make sure they are consistent. For example, two EMPIAR IDs: EMPIAR-10028 ^49^ and EMPIAR-10081 ^50^ with 300 micrographs (total 600 Cryo-EM micrographs) were used in cross-validation. The results of the 2D classes were compared based on total number of particles in each class, relative resolution of particles in the class, and distinct views of the structure of particles. Similar 2D classes, as shown in **Figure 12**, achieved by two independent Cryo-EM specialists validate the accuracy of the manually labelled particles.

**Figure 12:**
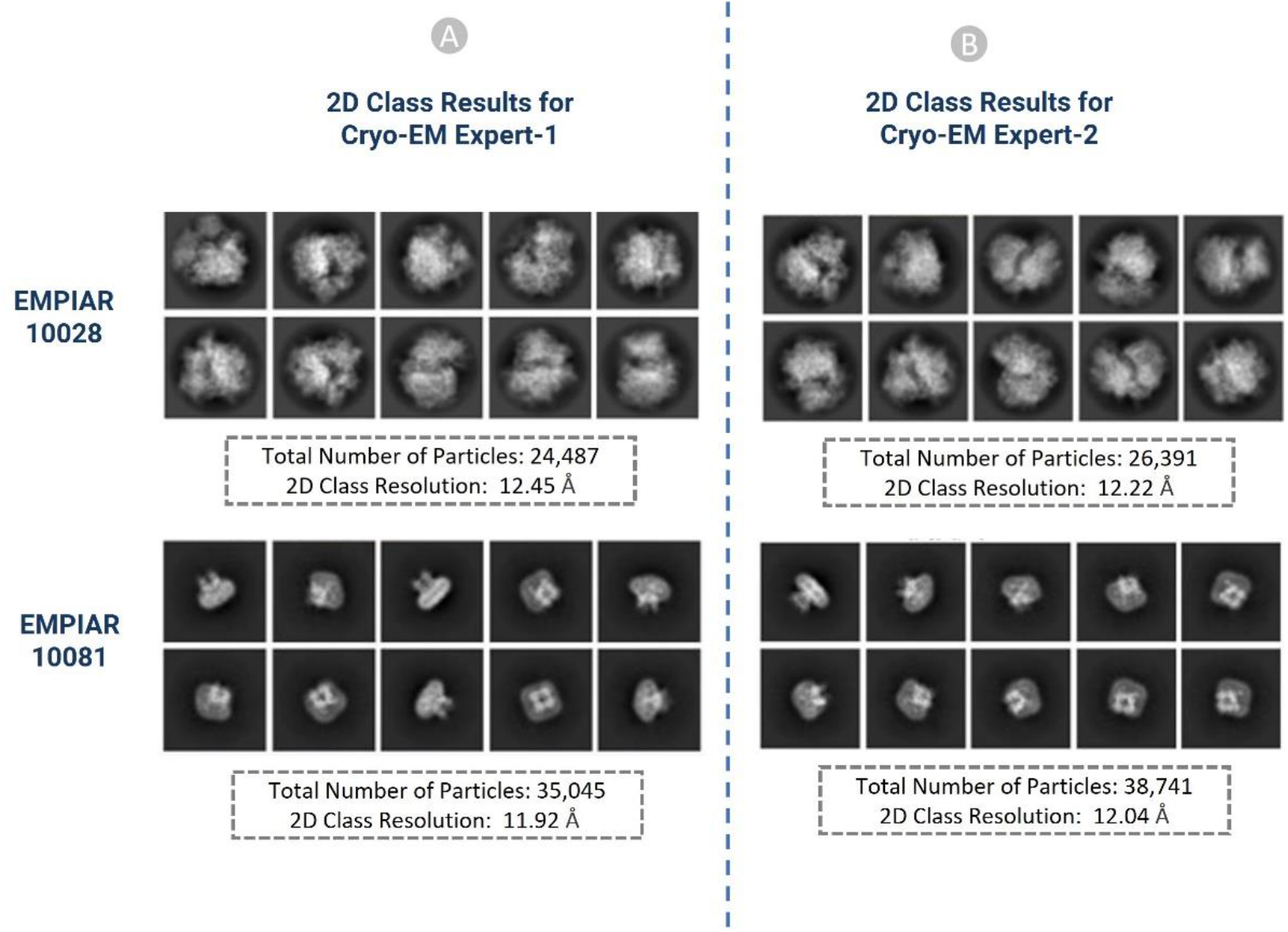
2D classification results of the picked particles of EMPIAR ID 10028 and 10081 (A): Results from Cryo-EM expert-1, (B): Results from Cryo-EM expert-2

### 4 Cross Validation with Gold Standard Particles Picked by the Authors

Gold standard particles are those particles that were picked by the Cryo-EM experts who generated the cryo-EM data. There are only a few EMPIAIR IDs deposited in EMPIAR that have both the micrographs and the gold standard particles. To validate the accuracy of our picked particles, we compared our results with the already-existing gold standard particles that are publicly available through the EMPIAR website. We carried out 2D and 3D validation for EMPIAR-10345 ^51^ and EMPIAR-10406 ^52^ to validate our particle labelling process as follows.

#### a) 2D Particle Class Validation with Gold Standard

In order to get the gold standard 2D particles of the dataset, we downloaded the particle stack image files (.mrc) and .star file with the attributes of picked particles from EMPIAR. We used the particle stack and the star files to create the 2D classification results using CryoSPARC. Eventually, we compared our 2D class results with the gold standard. We performed the comparison based on the total number of classes, total number of picked particles, resolution, and visual orientation of the protein particle for each EMPIAR ID. Our results and the gold standard results exhibit strong correlations. It is worth noting that a high number of particles alone does not necessarily yield high resolution. Selecting a decent number of high-quality particles spanning a wide angular distribution is important for generating high 2D and 3D resolution.

**Figure 13** shows the visual illustration 2D classification results for EMPIAR ID 10345 and EMPIAR ID 10406 published by the authors of the cryo-EM data and generated by us. They are consistent.

**Figure 13:**
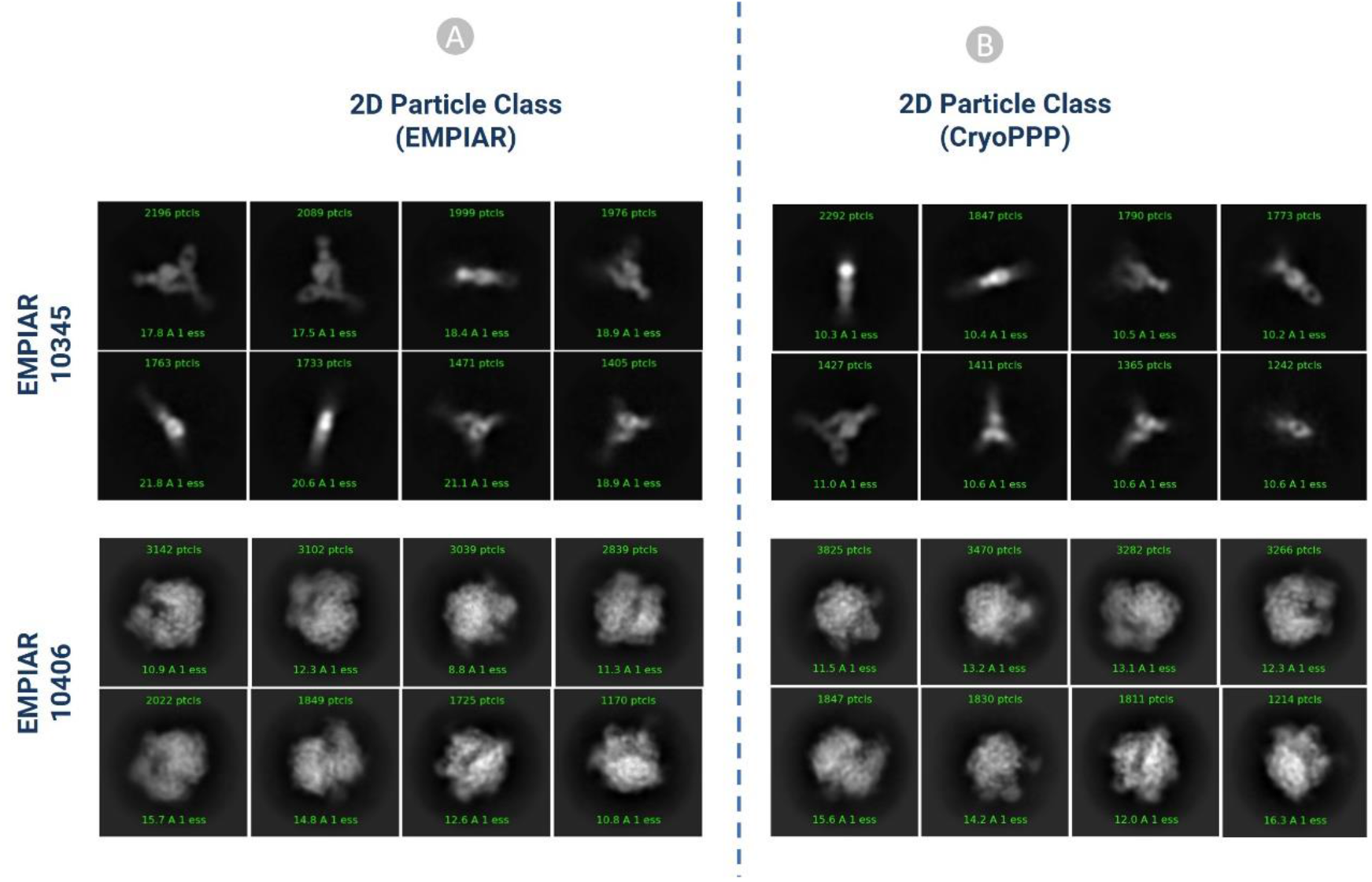
2D classification comparison for EMPIAR-10345 and EMPIAR-10406 (A) 2D classification published in EMPIAR. (B) 2D classification results of the particles by CryoPPP.

**Table 1** compares 2D classification results generated by authors and by us. In both cases, (**Figure 13(A)** and **Figure 13(B)**) the same 300 micrographs were used for comparison. On EMPIAR ID 10345, CryoPPP’s results have substantially higher resolution than the authors’ results for both N=50 and N=10 classes. On EMPIAR-10406, CryoPPP’s results have better resolution for N=50 particle classes and slightly lower resolution for N=10 particle classes.

**Table 1:** 2D classification result comparison for EMPIAR-10345 and EMPIAR-10406

| EMPIAR 10345 |  |  |
| --- | --- | --- |
|  | 2D Particle Class Statistics (EMPIAR) | 2D Particle Class Statistics (CryoPPP) |
| Number of Picked Particles | 17,838 | 15,894 |
| Weighted Average Resolution of 2D classes (N=50) | 18.63 Å | 10.25 Å |
| Weighted Average Resolution of 2D classes (N=10) | 20.52 Å | 10.53 Å |

| EMPAIR 10406 |  |  |
| --- | --- | --- |
|  | 2D Particle Class Statistics (EMPIAR) | 2D Particle Class Statistics (CryoPPP) |
| Number of Picked Particles | 23, 450 | 24,703 |
| Weighted Average Resolution of 2D classes (N=50) | 8.47 Å | 7.98 Å |
| Weighted Average Resolution of 2D classes (N=10) | 15.53 Å | 15.97 Å |

#### b) 3D Density Map Validation with Gold Standard

We performed an ab-initio reconstruction of the 3D density map using CryoPPP’s picked particles and compared the results with the gold standard 3D density maps from the EMPIAR website. The comparison of the 3D maps between EMPIAR and CryoPPP for EMPIAR-10345 and EMPIR-10406 is depicted in **Figure 14** and **Figure 15**. The results of 3D density maps, resolution, and direction distribution of protein particles are compared in the two figures.

**Figure 14:**
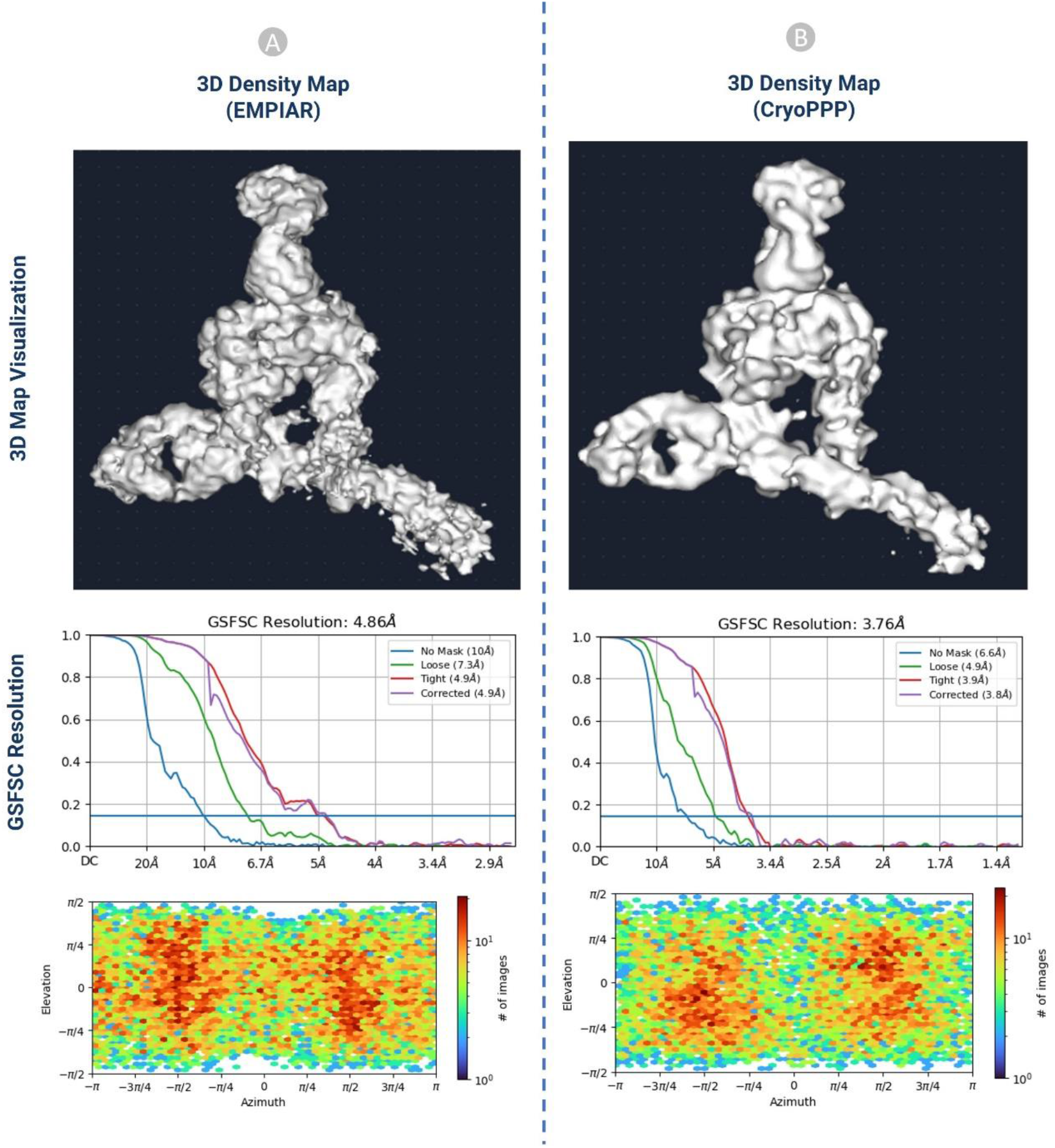
The comparison of 3D density maps, resolution, and direction distribution on EMPIAR-10345. (A) results published in EMPIAR. (B) results generated from the particles in CryoPPP.

**Figure 15:**
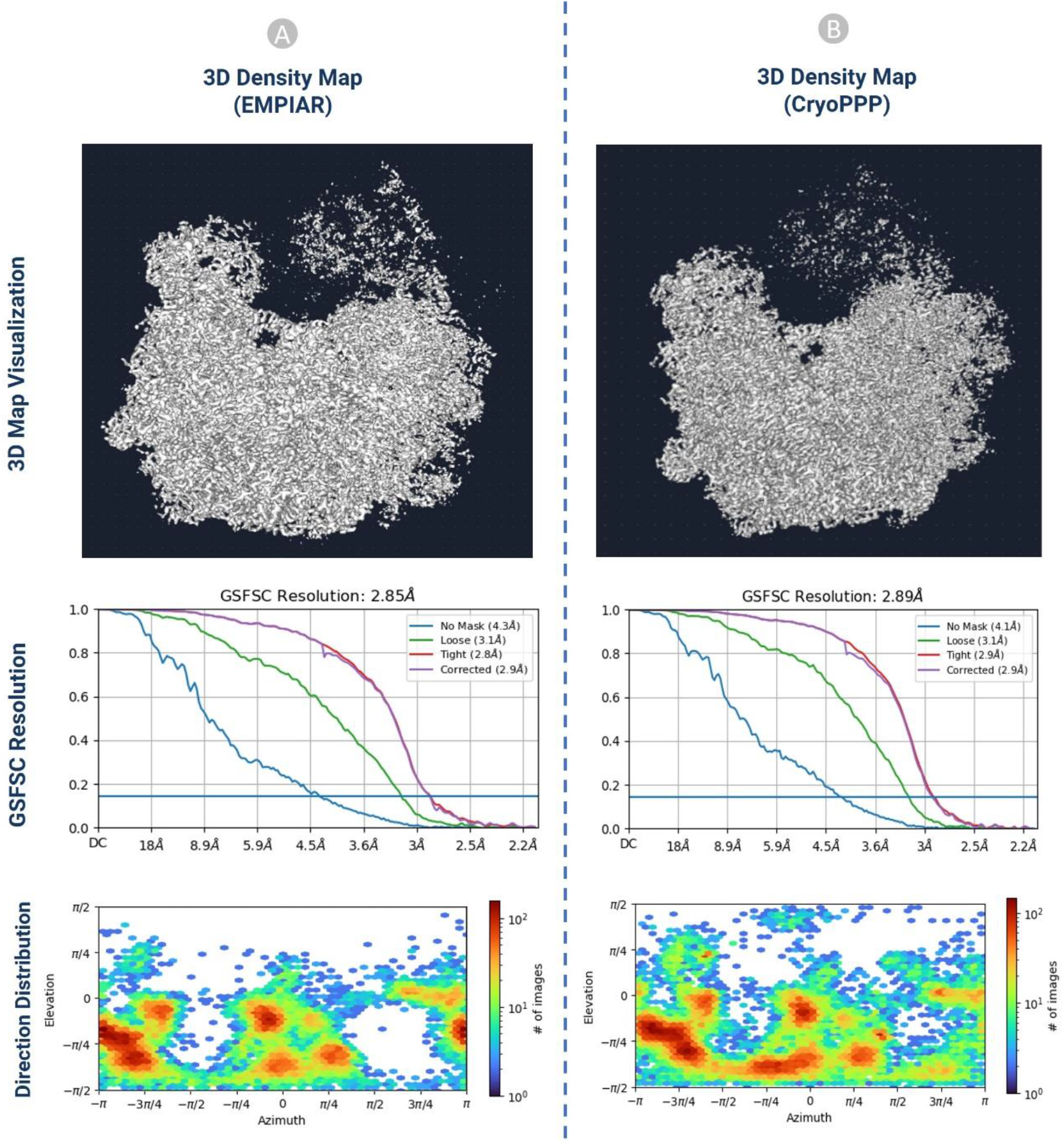
The comparison of 3D density maps, resolution, and direction distribution on EMPIAR-10406. (A) results published in EMPIAR. (B) results generated from the particles in CryoPPP.

The detailed comparison results are reported in **Table 2**. The ‘loose mask’ curve in the Fourier Shell Correlation (FSC) plots uses an automatically produced mask with a 15 Å falloff. The ‘tight mask’ curve employs an auto-generated mask with a falloff of 6 Å for all FSC plots. It is seen that CryoPPP outperforms in terms of all resolution (Gold Standard Fourier Shell Correlation (GSFSC), No mask, Loose, Tight and Corrected Mask) metrics on EMPIAR-10345 and achieved very similar results on EMPIAR-10406. This rigorous validation clearly demonstrates our manual particle picking procedure produced the high-quality picked particles in the CryoPPP dataset.

**Table 2:** 3D density map result comparison statistics for EMPIAR 10345 and EMPIAR 10406

| EMPIAR 10345 |  |  |
| --- | --- | --- |
|  | 3D Map Statistics (EMPIAR) | 3D Map Statistics (CryoPPP) |
| Number of Picked Particles | 17,838 | 15,894 |
| GSFSC Resolution | 4.86 Å | 3.76 Å |
| No Mask Resolution | 10 Å | 6.6 Å |
| Loose Mask Resolution | 7.3 Å | 4.9 Å |
| Tight Mask Resolution | 4.9 Å | 3.9 Å |
| Corrected Mask Resolution | 4.9 Å | 3.8 Å |

| EMPAIR 10406 |  |  |
| --- | --- | --- |
|  | 3D Map Statistics (EMPIAR) | 3D Map Statistics (CryoPPP) |
| Number of Picked Particles | 23, 450 | 24,703 |
| GSFSC Resolution | 2.85 Å | 2.89 Å |
| No Mask Resolution | 4.3 Å | 4.1 Å |
| Loose Mask Resolution | 3.1 Å | 3.1 Å |
| Tight Mask Resolution | 2.8 Å | 2.9 Å |
| Corrected Mask Resolution | 2.9 Å | 2.9 Å |

## Code Availability

The data analysis methods, software and associated parameters used in this study are described in the section of Methods. All the scripts associated with each step and the CryoPPP dataset are available at GitHub: https://github.com/BioinfoMachineLearning/cryoppp

## Acknowledgements

We thank the entire EMPIAR team for hosting the Cryo-EM data archive. Thanks to the researchers who deposited their cryo-EM images into EMPIAR for public use. We are grateful to Dr. Filiz Bunyak and Dr. Michael Chapman for their insights while curating CryoPPP.

## Author Contributions

J.C. conceived and conceptualized this research; A.D and R.G. designed the methodology; J.C and L.W. provided guidance through the experimental design; A.D. and R.G. wrote the scripts and codes to preprocess dataset; A.D and R.G. curated data; L.W. and J.C. conceptualized data validation; A.D. and R.G. drafted the manuscript; J.C and L.W. revised manuscript. All authors discussed the results, analyzed the data, and contributed to the final manuscript.

## References

1. Glaeser, R. M. Stroboscopic imaging of macromolecular complexes. Nat. Methods 10, 475–476 (2013).

2. Pakhrin, S. C., Shrestha, B., Adhikari, B. & Kc, D. B. Deep learning-based advances in protein structure prediction. Int. J. Mol. Sci. 22, (2021).

3. Boadu, F., Cao, H. & Cheng, J. Combining protein sequences and structures with transformers and equivariant graph neural networks to predict protein function. bioRxiv (2023) doi:10.1093/bioinformatics/xxxxx.

4. Dhakal, A., McKay, C., Tanner, J. J. & Cheng, J. Artificial intelligence in the prediction of protein-ligand interactions: recent advances and future directions. Briefings in Bioinformatics vol. 23 (2022).

5. Giri, N. & Cheng, J. Improving Protein–Ligand Interaction Modeling with cryo-EM Data, Templates, and Deep Learning in 2021 Ligand Model Challenge. Biomolecules 13, (2023).

6. Mahmud, S., Soltanikazemi, E., Boadu, F., Dhakal, A. & Cheng, J. Deep Learning Prediction of Severe Health Risks for Pediatric COVID-19 Patients with a Large Feature Set in 2021 BARDA Data Challenge. ArXiv (2022).

7. Grassucci, R. A., Taylor, D. J. & Frank, J. Preparation of macromolecular complexes for cryo-electron microscopy. Nat. Protoc. 2, 3239–3246 (2007).

8. Shen, P., Iwasa, J. & Brasch, J. Chapter 2: Cryo-EM grid preparation. https://cryoem101.org/chapter-2/ (2022).

9. Shen, P., Iwasa, J. & Brasch, J. Chapter 3: Grid Screening and Evaluation. https://cryoem101.org/chapter-3/ (2022).

10. Carragher, B. et al. Current outcomes when optimizing ‘standard’ sample preparation for single-particle cryo-EM. J. Microsc. 276, 39–45 (2019).

11. Chen, S. et al. High-resolution noise substitution to measure overfitting and validate resolution in 3D structure determination by single particle electron cryomicroscopy. Ultramicroscopy 135, 24–35 (2013).

12. Downing, K. H. & Hendrickson, F. M. Performance of a 2k CCD camera designed for electron crystallography at 400 kV. Ultramicroscopy 75, 215–233 (1999).

13. De Ruijter, W. J. Imaging properties and applications of slow-scan charge-coupled device cameras suitable for electron microscopy. Micron 26, 247–275 (1995).

14. Tang, G. et al. EMAN2: An extensible image processing suite for electron microscopy. J. Struct. Biol. 157, 38–46 (2007).

15. Scheres, S. H. W. RELION: Implementation of a Bayesian approach to cryo-EM structure determination. J. Struct. Biol. 180, 519–530 (2012).

16. Punjani, A., Rubinstein, J. L., Fleet, D. J. & Brubaker, M. A. CryoSPARC: Algorithms for rapid unsupervised cryo-EM structure determination. Nat. Methods 14, 290–296 (2017).

17. Marabini, R. et al. Xmipp: An image processing package for electron microscopy. J. Struct. Biol. 116, 237–240 (1996).

18. Heimowitz, A., Andén, J. & Singer, A. APPLE picker: Automatic particle picking, a low-effort cryo-EM framework. J. Struct. Biol. 204, 215–227 (2018).

19. Wang, F. et al. DeepPicker: A deep learning approach for fully automated particle picking in cryo-EM. J. Struct. Biol. 195, 325–336 (2016).

20. Zhu, Y., Ouyang, Q. & Mao, Y. A deep convolutional neural network approach to single-particle recognition in cryo-electron microscopy. BMC Bioinformatics 18, 1–10 (2017).

21. Xiao, Y. & Yang, G. A fast method for particle picking in cryo-electron micrographs based on fast R-CNN. AIP Conf. Proc. 1836, (2017).

22. Wagner, T. et al. SPHIRE-crYOLO is a fast and accurate fully automated particle picker for cryo-EM. Commun. Biol. 2, 1–13 (2019).

23. Zhang, J. et al. PIXER: An automated particle-selection method based on segmentation using a deep neural network. BMC Bioinformatics 20, 1–14 (2019).

24. Yao, R., Qian, J. & Huang, Q. Deep-learning with synthetic data enables automated picking of cryo-EM particle images of biological macromolecules. Bioinformatics 36, 1252–1259 (2020).

25. Tegunov, D. & Cramer, P. Real-time cryo-electron microscopy data preprocessing with Warp. Nat. Methods 16, 1146–1152 (2019).

26. Bepler, T. et al. Positive-unlabeled convolutional neural networks for particle picking in cryo-electron micrographs. Nat. Methods 16, 1153–1160 (2019).

27. Al-Azzawi, A., Ouadou, A., Tanner, J. J. & Cheng, J. AutoCryoPicker: an unsupervised learning approach for fully automated single particle picking in Cryo-EM images. BMC Bioinformatics 20, 326 (2019).

28. Al-Azzawi, A. et al. DeepCryoPicker: fully automated deep neural network for single protein particle picking in cryo-EM. BMC Bioinformatics 21, 1–38 (2020).

29. Iudin, A. et al. EMPIAR: the Electron Microscopy Public Image Archive. Nucleic Acids Res. 51, D1503–D1511 (2023).

30. Agard, D., Cheng, Y., Glaeser, R. M. & Subramaniam, S. Single-particle cryo-electron microscopy (cryo-EM): Progress, challenges, and perspectives for further improvement. Advances in Imaging and Electron Physics vol. 185 (Elsevier, 2014).

31. Langlois, R. et al. Automated particle picking for low-contrast macromolecules in cryo-electron microscopy. J. Struct. Biol. 186, 1–7 (2014).

32. Baldwin, P. R. & Penczek, P. A. The Transform Class in SPARX and EMAN2. J. Struct. Biol. 157, 250–261 (2007).

33. Zhang, C. et al. TransPicker: A Transformer-based Framework for Particle Picking in cryoEM Micrographs. Proc. - 2021 IEEE Int. Conf. Bioinforma. Biomed. BIBM 2021 1179–1184 (2021) doi:10.1109/BIBM52615.2021.9669524.

34. George, B. et al. CASSPER is a semantic segmentation-based particle picking algorithm for single-particle cryo-electron microscopy. Commun. Biol. 4, 1–12 (2021).

35. McSweeney, D. M., McSweeney, S. M. & Liu, Q. A self-supervised workflow for particle picking in cryo-EM. IUCrJ 7, 719–727 (2020).

36. Azzawi, A. Al, Ouadou, A., Tanner, J. J. & Cheng, J. A super-clustering approach for fully automated single particle picking in cryo-em. Genes (Basel). 10, (2019).

37. Mallick, S. P., Zhu, Y. & Kriegman, D. Detecting particles in cryo-EM micrographs using learned features. J. Struct. Biol. 145, 52–62 (2004).

38. Hoang, T. V., Cavin, X., Schultz, P. & Ritchie, D. W. GEMpicker: A highly parallel GPU-accelerated particle picking tool for cryo-electron microscopy. BMC Struct. Biol. 13, (2013).

39. Wagner, T. & Raunser, S. The evolution of SPHIRE-crYOLO particle picking and its application in automated cryo-EM processing workflows. Commun. Biol. 3, 1–5 (2020).

40. Masoumzadeh, A. & Brubaker, M. HydraPicker: Fully automated particle picking in cryo-em by utilizing dataset bias in single shot detection. 30th Br. Mach. Vis. Conf. 2019, BMVC 2019 (2020).

41. Campbell, M. G. et al. Movies of ice-embedded particles enhance resolution in electron cryo-microscopy. Structure 20, 1823–1828 (2012).

42. Rawson, S., Iadanza, M. G., Ranson, N. A. & Muench, S. P. Methods to account for movement and flexibility in cryo-EM data processing. Methods 100, 35–41 (2016).

43. Noble, A. J. et al. Routine single particle CryoEM sample and grid characterization by tomography. Elife 7, 1–42 (2018).

44. Singer, A. & Sigworth, F. J. Computational Methods for Single-Particle Cryo-EM. 1–40 (2020).

45. Li, J. et al. Cryo-EM structures of Escherichia coli cytochrome bo3 reveal bound phospholipids and ubiquinone-8 in a dynamic substrate binding site. Proc. Natl. Acad. Sci. U. S. A. 118, (2021).

46. Scheres, S. H. W. Semi-automated selection of cryo-EM particles in RELION-1.3. J. Struct. Biol. 189, 114–122 (2015).

47. Pettersen, E. F. et al. UCSF Chimera - A visualization system for exploratory research and analysis. J. Comput. Chem. 25, 1605–1612 (2004).

48. Pettersen, E. F. et al. UCSF ChimeraX: Structure visualization for researchers, educators, and developers. Protein Sci. 30, 70–82 (2021).

49. Wong, W. et al. Cryo-EM structure of the Plasmodium falciparum 80S ribosome bound to the anti-protozoan drug emetine. Elife 2014, 1–20 (2014).

50. Lee, C. H. & MacKinnon, R. Structures of the Human HCN1 Hyperpolarization-Activated Channel. Cell 168, 111-120.e11 (2017).

51. Campbell, M. G. et al. Cryo-EM Reveals Integrin-Mediated TGF-b Activation without Release from Latent TGF-b Article Cryo-EM Reveals Integrin-Mediated TGF-b Activation without Release from Latent TGF-b. Cell 180, 490-501.e16 (2020).

52. Nicholson, D., Edwards, T. A., O’Neill, A. J. & Ranson, N. A. Structure of the 70S Ribosome from the Human Pathogen Acinetobacter baumannii in Complex with Clinically Relevant Antibiotics. Structure 28, 1087-1100.e3 (2020).

